# Case based learning as a dynamic approach towards learning oral pathology

**DOI:** 10.1101/2021.04.02.438062

**Authors:** Ashish Shrestha, Vinay Marla, Jyotsna Rimal, Sushmita Shrestha, Shashi Keshwar, Jia Zhimin

**Affiliations:** Department of Oral Pathology, BP Koirala Institute of Health Sciences, Dharan, Nepal; Department of Oral Pathology, Penang International Dental College, Penang, Malaysia; Department of Oral Medicine and Radiology, BP Koirala Institute of Health Sciences, Dharan, Nepal; Department of Conservative Dentistry and Endodontics, BP Koirala Institute of Health Sciences, Dharan, Nepal; Director of SMU- FAIMER Regional Institute and Deputy Dean, Shenzhen Hospital of Southern Medical University, Guangzhou, China

**Keywords:** case-based learning, dental students, learning environment, oral pathology

## Abstract

**Introduction:** Oral pathology comprises of extensive teaching-learning of histopathology. Teaching histopathology is always challenging more when numerous histopathology slides are involved. There is lack of interest among the students and also face challenges to correlate with the clinical presentation. For effective learning, need for a method which is interactive and helps create clinico pathological correlation was felt. This study was designed to assess the impact and feasibility of case-based learning in routine teaching among dental students.

**Methods:** A cross sectional study was conducted among 58 undergraduate dental students, wherein a case-based learning method was introduced in the practical classes. Thirty students were randomly selected as case based learning and remaining as conventional group. Multiple paper-based cases on oral cancer was designed and used for the case based learning group of students. A self-designed pre and posttest interventional tools along with post intervention assessment using modified essay question was designed and applied.

**Results:** Significant difference in the scores between pre and post intervention questionnaire was observed among the case-based learning group (p<0.0001), however non-significant among the conventional. Similarly, significant difference in the scores of modified essay question was observed between the study groups. Majority of the students among the case-based learning group agreed that it benefitted their personal, professional and communicative skills. The students expressed their enthusiasm in learning with this method and suggested to apply it more often.

**Discussion:** Case-based learning is an effective method and incorporation of it on a regular basis could help favor an effective learning environment.

## Introduction

Histopathology has been an important component of curriculum of any medical or dental school. Apart from theoretical knowledge about histopathology, the practical sessions are equally important. The practical sessions provide students with intuitive perceptions of the pathologic basis of disease to either observe the histopathology slides directly under the microscope or digitally [1]. However, the conceptual difficulty and lack of interest, due to its supposed irrelevance to their future clinical or professional activities, render students lacking the motivation to learn histopathology. Students tend to take the subject seriously only if they pursue it as a specialty subject for their higher education. Moreover, the conventional practices of viewing fixed focus of the histopathology slides, make it dreary and tedious to the students; and at times disconnects them from the subject. Hence they fail to realize the connection between clinical presentations and histopathological features.

Case based learning (CBL) has been receiving attention in medical education as it is a student-centered teaching methodology that exposes students to real-world scenarios that need to be solved using their reasoning skills with existing theoretical knowledge [2]. The cases in case based learning situates the information in real world contexts. Contextualization enhances learning by providing association that facilitates memory storage, retention, and retrieval, thereby promoting motivation towards learning [3]. Students can acquire adequate knowledge about patient care by accessing real cases. They can gain better understanding of various individual perspectives, which is useful for developing their mindset for cooperation, as well as continuous knowledge development [4].

In context to oral pathology practical sessions, CBL might orient students to correlate clinical details with the histological picture thereby increasing their cognitive and problem solving ability and also helping them realize its significance. Incorporation of CBL could raise students’ interest in the subject and motivate them towards active learning and improve academic performance [5– 7]. In the present study, we have applied the concept of CBL in practical session of oral pathology with an aim to introduce CBL in Oral Pathology in our institution and assess its impact and feasibility.

## Materials and methods

### Study design and study population

A cross sectional study was conducted among the third year undergraduate dental students in the department of Oral Pathology. Among the total of 59 students, 58 participated in the study. The students were divided into: Group I (case based learning group) and Group II (conventional group) after obtaining consent. The study was conducted after obtaining ethical clearance from the Institute Review Committee.

### Study setting

The study was conducted during the regular laboratory exercise (LABEX) of the department. No prior CBL session had been conducted in the department.

### The teaching approach

A paper based Case Based Learning session along with video recordings was designed. The topic of ‘oral squamous cell carcinoma’ (OSCC) was decided by the panel of faculties from the department. Prior to the intervention, the theory class on OSCC was conducted as structured interactive session (SIS). Two days’ study module, each of three hours, was designed to conduct the practical session (Figure 1).

**Figure 1:**
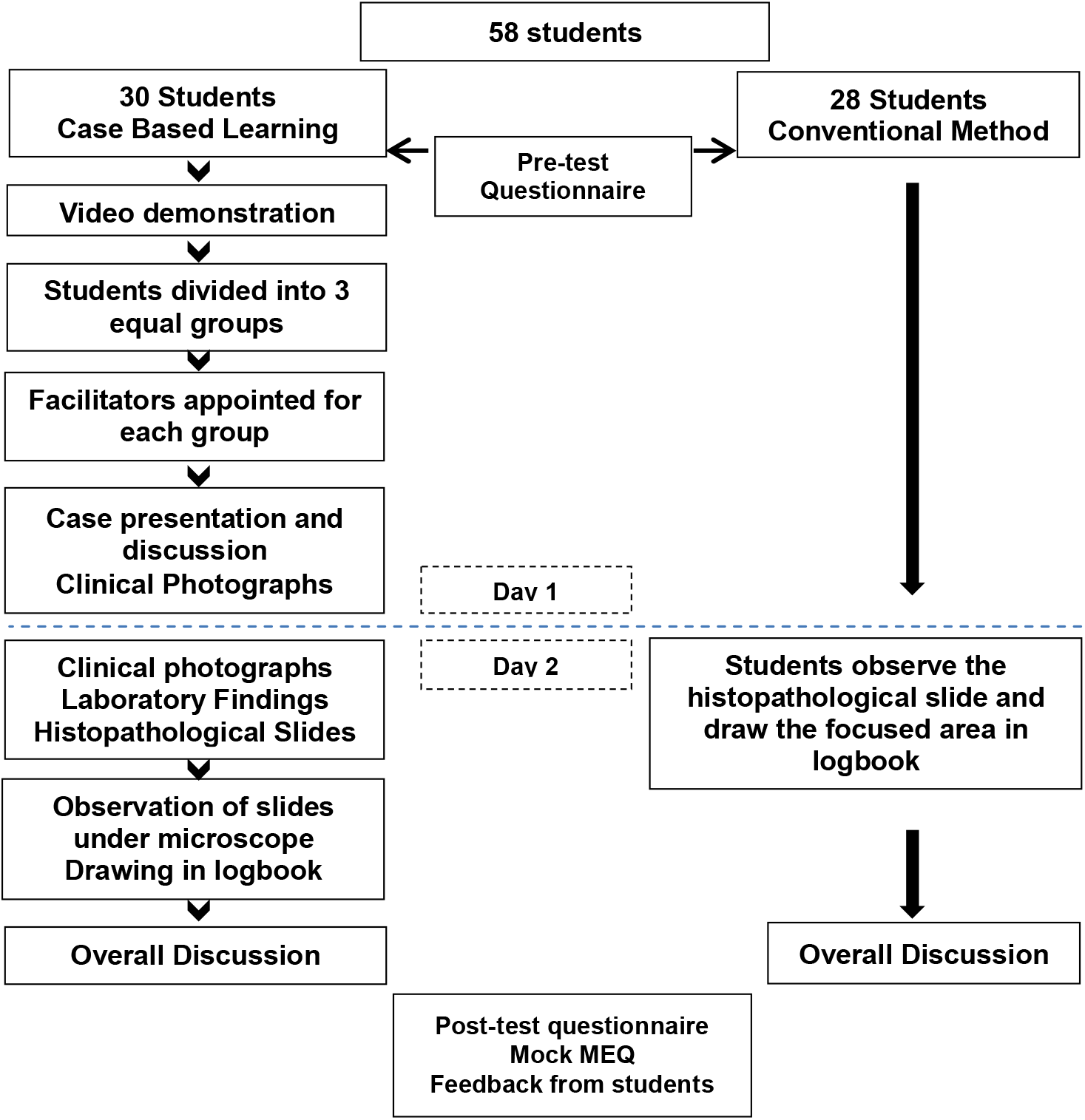
Schematic outline of the study details

The students were randomly divided into CBL group (30 students) and conventional learning group (28 students). The CBL groups were further divided into three equal groups and individual facilitators were assigned to each. Three residents from the College of Dental Surgery were requested to take the role of facilitators for each sub-group. All the facilitators were trained and provided with a facilitators guide to keep the learning standard among three groups. After the completion of the study, students among the conventional group underwent the same CBL session; however the results were not used for statistical evaluation.

### Tools for content and process validation

a. *Case selection and design* A session on ‘oral squamous cell carcinoma’ was finalized with five cases of OSCC and the learning objectives were defined as per the curriculum. The cases had varying clinical presentations and histopathological grades. Detailed clinical history, habit history, lesion descriptions, radiographs and laboratory investigations, along with final histopathological diagnosis were included. Colored photographs of the lesions, histopathological features and printed images of the radiographs were also prepared. Prior consent had been obtained from the patients to use the relevant information and photographs without disclosing the personal information, as per the routine protocol of the department.
b. *Video preparation* An educational video (6.51 minutes) with information regarding the oral cancer, biopsy and tissue processing, and slide preparation was prepared by the department. The video also comprised of a special segment drawing in the experiences of the oral cancer patients.
c. *Facilitators guidelines* A guideline was prepared by the faculties of the department and the members of Health Professions Education Department of the Institute. Structured instructions regarding CBL process, learning objectives and possible outcomes of each sub-session, time frame, feedback of students, etc. were included.
d. *Pre-test and post-test questions* Multiple choice questions (MCQs) in accordance with the learning objectives of the topic were prepared by the faculties of the department. The self-designed MCQ consisting of 19 items were given to 20 students from fourth year as a pilot study. The reliability of the instrument was assessed using Cronbach’s alpha which showed better result with the coefficient 0.723 after excluding three items. The intra correlation coefficient was estimated to be 0.723 (95% CI; 0.508 −0.872). The final scale included only 16 items and was used for assessing the knowledge on OSCC and for the comparison of pre and post intervention (CBL) scores. The questions were scenario-based and included information regarding background, epidemiology, clinical presentations, radiographic and laboratory findings, histopathology and prognosis of OSCC. Each MCQ had one best answer with three distractors. Each question was allocated one mark for correct response.
e. *Mock Modified Essay Questions (MEQ)* A mock MEQ (20 marks) was designed by the faculties of the department, divided into eight sub-questions. The sub set of MEQ was sequentially given, only after completion of the previous set.
f. *Students feedback forms* A seventeen items evaluation form was adapted and modified to assess the personal (10 questions), professional (four questions) and communicative (three questions) benefits from the case based learning method [8, 9]. The sessions were conducted simultaneously between the CBL and conventional group in different rooms.

### Statistical analysis

The data were entered in Microsoft Excel spreadsheets and transferred to SPSS version 11.5 for further statistical analysis. The quantitative data were analyzed using appropriate statistical tool. The statistical analyses were considered as significant at p value less than 0.05.

## Results

Of the total (59 students), 58 students provided consent to participate in the study. There were six females and 24 males in the CBL group while 17 females and 11 males in the conventional group. All students were within an age range of 19 to 23 years.

No significant differences were found, when comparing the pretest scores of CBL and conventional group (p=0.109) and comparing the pre-test and post-test scores of the conventional group (p=0.44). However, a highly significant difference was observed in the pretest and posttest comparison of the CBL group (p<0.0001) and in between CBL and conventional group (p=0.0001). The score range was 8-14 for CBL and 3-12 for conventional group. Fourteen of the conventional group had score below eight. A significant difference in the mean scores of modified essay question was observed between the conventional and CBL groups (p=0.001). The maximum score among CBL group students was 18.5 and minimum of 13, however the maximum was 14 and minimum of 4 among the conventional group.Most of the students in their feedbacks said that learning about OSCC by case-based learning was interesting and helpful. Majority of the students agreed that they benefitted professionally and personally moreover they also improved their communication skills (Table 1). Almost 80% of the students strongly agreed that CBL was better than conventional method. At the end of the session Group I students were asked to note down their experiences regarding CBL with an open feedback which were subsequently categorized (Table 2). Feedback from the facilitators was promising and encouraging. They expressed that the students were motivated to learn with this new method, which could be an important learning experience for them.

**Table 1:** Students responses towards case based learning

| **Students perceived professional benefits** | **Strongly Agree** | **Moderately Agree** | **Neutral** |
| --- | --- | --- | --- |
| • CBL stimulated my desire to learn | 15 | 15 | - |
| • I feel confident to apply basic sciences and oral pathology concepts to solve clinical cases | 14 | 13 | 3 |
| • CBL is good method to practice integration of knowledge and skill | 24 | 6 | - |
| • CBL improved my clinical reasoning ability | 14 | 16 | - |
| • I was motivated to learn oral pathology by CBL | 6 | 22 | 2 |
| • The emphasis on clinical concept was detrimental to learn oral pathology | 8 | 20 | 2 |
| • The CBL helped to reinforce concepts taught in class | 17 | 13 | - |
| • CBL promoted self-directed learning skills | 14 | 16 | - |
| • CBL has increased my self-confidence and attitude towards learning | 13 | 13 | 4 |
| • CBL is better than traditional teaching method | 24 | 6 | - |
| **Students perceived communication benefit** |  |  |  |
| • I understand what makes small groups operate effectively | 18 | 12 | - |
| • I am confident in my ability to contribute effectively to the team | 17 | 12 | 1 |
| • I am confident in discussing my ideas with my teammates | 14 | 15 | 1 |
| • I am confident in sharing my ideas with my facilitators | 13 | 16 | 1 |
| **Students perceived personal benefit** |  |  |  |
| • My participation in this course was personally enjoyable | 17 | 12 | 1 |
| • My participation in this course positively impacted my personal relationships with colleague and facilitators | 8 | 16 | 6 |
| • My participation in this course positively impacted my professional relationships | 15 | 13 | 2 |

**Table 2:** Categorization of feedback from the students

| Category | Students responses |
| --- | --- |
| **Learning environment** | • Interactive, fun learning, easy understanding in small group<br>• Give 'equal chance' to everyone to open up with their opinion<br>• Innovative learning, help us to brainstorm<br>• The idea of video, slide and image were helpful<br>• One-to-one interaction with mentor help us to put our views properly |
| **Organization and design** | • Real case video was very effective, feeling of real scenario<br>• CBL gave real situation of diseases, taught the way to approach<br>• Topic were divided into sub-topic hence discussion was easy for knowledge sharing<br>• Liked the concept of video but the sound quality was not up to mark |
| **Acquisition of skill and knowledge** | • Help to gain idea about clinical approach to the case and to know about various screening test done in clinical practice<br>• Best method of learning that improved my clinical reasoning ability<br>• More retentive and more helpful |
| **Personal development** | • Improved our team work and relationship with friends<br>• Encourage the student who speak<br>• Motivate toward the self-learning<br>• It increased self-confidence and attitude towards learning |
| **Way forward** | • This would be fruitful if conducted with most of the commonly encountered cases in clinic<br>• Should be done more often for important topics that go neglected by just theoretical knowledge<br>• CBL should be arranged every month. If this had been conducted earlier it would have been more beneficial.<br>• Should be conducted in patient itself also |

## Discussion

Case based learning module on ‘oral squamous cell carcinoma’ was designed and applied on third year undergraduate dental students. OSCC was chosen as it has a significant importance in dental curriculum, and is one of the commonly encountered neoplasms not only in locally in Nepal but also globally [10].

Existing knowledge on the topic was analyzed by comparing pre-intervention scores using 16-items questionnaire, where no statistical differences between the study groups were observed. This was an ideal scenario since it allowed better comparison of the students’ scores. But there was heterogeneous distribution of students according to gender. Following the intervention, same questionnaire was used, wherein the CBL group scored higher (p<0.001) compared to their pre-intervention scores. However, the conventional group showed only marginal improvement. Further comparison between the CBL group and the conventional group showed statistically significant differences (p=0.001), suggesting better performance by the CBL group students. Similar findings were observed in a study conducted in China by Du G et al, who implemented CBL for Oral Medicine among fourth year dental students and compared it to lecture-based education. They found that the students with CBL scored higher when compared to the students receiving lecture-based education [11]. Another study conducted among first year dental students for physiology also yielded improvement of scores following CBL sessions [12]. Likewise, CBL also led to improvement of test scores among a cohort of students in Implant Dentistry [13]. The improvement in scores in our study might be attributed to the dynamic methods of learning in CBL in Group I students compared to the conventional methods in Group II, which predominantly relied on existing theoretical knowledge and memorization of histopathological images. In the traditional method, students need an inherent self-directed active learning for better performance, and the majority of students end up with an average performance. According to Hansen and Krackov, CBL promoted active discussion and participation [14]. Also, Sutyak et al stated that through CBL, students engaged in problem solving activities, seeking support and feedback from colleagues and experts [15].

Eighty percent (n=24) of the students strongly agreed that CBL was better and further strongly agreed that it was a good method to practice integration of knowledge and skill. Majority of students (n=17) also strongly agreed that CBL helped them reinforce the concepts. Few students chose to remain neutral on questions involving their self-confidence, attitude and motivation towards learning. This could be due to the inclination of students towards the clinical subjects as opposed to Oral Pathology as their choice of subject in dentistry. Majority of the students accepted that small group discussions were effective. They asserted that they felt confident to contribute to the team and also enjoyed the learning process. Overall, the student’s feedback suggested that it was very well accepted and helped in improving their learning capabilities.

Majority of the studies conducted worldwide among dental students and others have given positive feedback for CBL and preferred it over traditional methods [8, 16–18] suggesting incorporation of CBL in the curriculum. Research conducted on a CBL based curriculum in a dental school in Germany involving 404 participants suggested that the students perceived CBL to benefit their research competence, interdisciplinary thinking, dental-medical knowledge, practical dental skills, team work and independent learning [16]. Botelho and O’Donnel in their study on fixed Prosthodontic simulation laboratory course found that students valued aspects of small group discussion, the most. The students were also able to better correlate the topic clinically as opposed to learning through lecture. They also suggested that small group discussions allowed integration of information from different sources, utilizing broader concepts to specific cases [19]. Improvement in perceived satisfaction, motivation and engagement among students was reported when CBL was incorporated in a Pharmacology course [20]. In a study conducted in dental materials, large number of students reported an increase in motivation to learn with the CBL method [17]. According to Steinhert, the students highlighted the role of group atmosphere, facilitation skills and clinical relevance of the cases in effective small group leaning during CBL [21].

The students considered CBL as the preferred method of learning over the existing method. The existing method involves background theoretical knowledge along with observation of static microscopic fields. Oral pathology is best learned on the principles of clinico-pathological correlation. The traditional method creates a lacuna in the understanding, due to which the students are not able to effectively understand the pathologic basis of the disease by just observing the slides. CBL is an active learning process which offers the advantage of simulation of clinical cases helping them to understand the nature of disease before evaluating the histopathology of the case. This approach can also lead to increased retention of information and better performance in assessments [5]. According to a study conducted among first year dental students, majority of the students felt that CBL provoked interest to learn and was effective in linking basic concept with clinical application. Majority felt that it promoted critical thinking, clinical reasoning, problem solving and indicated their preference of CBL for other topics as well [12].

The attempt of CBL has been very rewarding with this study as evident by the feedback from the students, who encouraged it to be conducted regularly. Irrespective of the mode of teaching histopathology, either digitally or by direct viewing under microscope; incorporation of innovative learning and assessment methods has always been motivating. A proper lesion plan and class room environment can impart good learning outcome. The existing curriculum of BP Koirala Institute of Health Sciences, Nepal has been designed following the SPICES model (student-centered, problem-based, integrated, community based, electives, and systematic) which encourages early clinical exposure, students centered and multidisciplinary learning approach [22]. Oral pathology is a bridge between the basic sciences and clinical dentistry, and adaptation of case based learning could help student to acclimatize and prepare for subsequent academic years which could enhance the learning outcome and overall performance of the students.

## Conclusion

Case-based learning is an effective teaching-learning method especially in professional education courses like dentistry. The present study suggests positive impact in learning oral pathology by incorporation of case-based learning on a regular basis. Moreover it could also be an example for other disciplines to adapt this innovative teaching-learning method.

## Acknowledgements

We would like to thank all our students for participating in the study.

